# Neuromodulation of pelvic nerve during early phase of spinal cord injury in rats using implantable prototype devices - a preliminary study

**DOI:** 10.1101/759449

**Authors:** Wendy Yen Xian Peh, Monzurul Alam, Rakib Uddin Ahmed, Sanghoon Lee, Li Jing Ong, Shuhao Lu, Roshini Mogan, Gil Gerald Lasam Gammad, Kian Ann Ng, Astrid Rusly, Sudip Nag, Yong-Ping Zheng, Shih-Cheng Yen

**Author notes:** **Corresponding author**: Dr. Shih-Cheng Yen.

## Abstract

**Objectives:** The bladder becomes retentive during the early phase of spinal cord injury, and requires proper bladder management to prevent damage to the lower urinary tract and kidney. We investigated the effects of on-demand pelvic nerve stimulation on the areflexive bladder during the earliest phase of complete spinal cord injury in rats and the use of pelvic nerve signals as a proxy to estimate intravesical pressure for closed-loop applications.

**Materials and Methods:** In order to stimulate the pelvic nerves in female Sprague-Dawley rats with complete spinal cord transection (T7 level), a flexible electrode was implanted unilaterally on pelvic nerve, and electrical stimulation was provided by a custom-built nerve stimulator. Stimulation-evoked voiding was monitored in the awake state while size, capacity and spontaneous contractions of the bladder were analysed under anaesthesia. Separately, recordings of the pelvic nerve signals, external urethral sphincter activity and intravesical pressure were performed in animals with intact and transected spinal cord under anaesthesia.

**Results:** Successful pelvic nerve stimulation enabled more frequent voiding, reduced overdistension of bladder, and preserved non-voiding spontaneous bladder contractions. Typical bladder management protocol for SCI rats (manual expression every 8 – 12 hours) resulted in more severe bladder overdistention. Signal processing of the recorded extraneural pelvic nerve signals successfully reconstructed changes in intravesical pressure, demonstrating their use in estimating the fullness and contractions of the bladder.

**Conclusions:** The preliminary results suggest that pelvic nerve stimulators can serve as an alternative method for frequent emptying of the areflexive bladder. Simultaneous recording of the same pelvic nerve will be useful for development of a closed-loop neuroprosthesis.

## 1. Introduction

Spinal cord injury (SCI) can disrupt the communication of sensory and parasympathetic efferent circuits at the lumbar and sacral levels with that of supraspinal brain regions required for the perception of bladder fullness and voluntary control of micturition^1^. Complete SCI results in an initial spinal shock phase whereby the bladder becomes areflexive and retentive^2^. For human patients with the inability to empty the bladder sufficiently, indwelling or intermittent catheterization remain the primary bladder management techniques. However, catheterization suffers from potential complications such as urinary tract infections^3^. While early sacral neuromodulation in complete SCI patients prevented the development of urinary incontinence, the patients were not able to void on demand^4^. A possible alternative to restore voiding in the areflexive bladder is to activate the parasympathetic pathway for bladder contraction, either by stimulating the pelvic nerve or efferent nerve endings in the bladder wall^5–9^, or in the case of complete SCI patients, sacral anterior root stimulation (SARS) by the commercially available Finetech-Brindley device^10^. However, SARS is usually available as a treatment only many months after injury, and requires a dorsal rhizotomy^11^. Therefore, not much is understood about the usefulness of SARS in the acute stages of SCI. Neuromodulation studies for bladder dysfunction in animal models of SCI are also mostly focused on the chronic stages of SCI where bladder hypertrophy, overactive bladder, and detrusor-sphincter dyssynergia are observed^12–15^. These animal studies typically used manual expression (or the Credé maneuver) twice to three times daily, whereby pressure is applied gently to the abdominal area over the bladder to overcome urethral pressure to enable urine release, for early post-operative bladder management. Thus, more studies are needed to understand the early effects of electrical stimulation of the efferent parasympathetic pathway on the areflexive bladder.

In this preliminary study, we focused on investigating the effects of pelvic nerve stimulation on the areflexive bladder during the acute phase (within 28 hours) of the complete thoracic cord injury in awake rats using implantable flexible electrode and a prototype stimulator. The pelvic nerve branches off from the L6-S1 nerve trunk, and contains axons of preganglionic parasympathetic neurons that synapse onto the postganglionic parasympathetic neurons in the major pelvic ganglion in the rodents, which then release acetylcholine to excite the smooth detrusor muscles in the bladder wall^1^. In order to perform nerve stimulation in awake SCI rats, we implanted a bipolar flexible neural clip^16,17^ on the pelvic nerve, and used a prototype of a wireless nerve stimulator to deliver biphasic constant current pulses. Bladder size, capacity, and preservation of non-voiding spontaneous contractions^18^ were monitored and quantified. As the pelvic nerve also carries sensory afferents that convey information from mechanoreceptors and nociceptors within the bladder wall^1^, we also performed extraneural recordings from pelvic nerve using both commercial benchtop system as well as implantable neural amplifier to examine the possibility of using pelvic nerve signals as a proxy to estimate intravesical pressure changes for closed-loop applications^19^.

## 2. Materials and Methods

### 2.1 Animal Subjects

15 adult female Sprague-Dawley (SD) rats (245-300 g body weight) were used in this study. Experimental procedures were performed in accordance with the guidelines and approval of the Animal Subjects Ethics Sub-committee of the Hong Kong Polytechnic University, and the Institutional Animal Care and Use Committee of the National University of Singapore.

### 2.2 Surgery for pelvic nerve stimulation in SCI rats

Each subject was anesthetized with isoflurane and the body temperature was maintained at 37 °C. The surgical sites were shaved and disinfected by povidone-iodine followed by 70% ethanol, and the animal was placed in the prone position with its head secured to a stereotaxic frame. In order to carry out a complete spinal cord injury (SCI) at the T7 level, a dorsal midline incision was carried out at the T5-T9 level. Fascia and muscles over the T7 vertebra were removed to isolate the spinous process with rongeurs, and the dura was opened by using a sharp needle. The cord was then cut by using a fine micro spring scissors to produce a complete transaction. After the SCI, skin incisions were made at the abdominal and head area to allow the head-plug, with connecting wires (DFT^™^ wire, Fort Wayne Metals) to the flexible neural clip electrodes coated with iridium oxide for the pelvic nerve^16^ and the ground electrode (Cooner Wire AS632), to be tunnelled subcutaneously from the abdominal skin incision towards the skull for implantation. Metal screws and dental cement were used for anchoring the head-plug on the rat’s skull. The rat was then removed from the stereotaxic frame, and turned to the supine position for an abdominal muscle incision to expose the left pelvic nerve branches as previously described^9^. The electrode was then implanted onto the exposed pelvic nerve, and secured with silicone elastomer (Kwik-Sil, World Precision Instruments). At the end of the surgery, all the exposed muscles, connective tissue, and the skin were sutured by using 4.0 Ethicon sutures. Post-surgery oral antibiotic (Enrofloxacin 0.6 ml/100 ml water) and analgesic (Buprenorphine HCL 0.5mg/kg, S.C.) were administered.

### 2.3 Electrical stimulation of pelvic nerve in awake SCI rats

The nerve stimulator prototype used in this study was modified from a custom-built wireless muscle stimulator system^20^, to deliver biphasic rectangular constant current pulses at an amplitude of 400 μA, 20 Hz frequency, phase width 300 μs, and lasting 10 s per train. A graphic user interface on a laptop was used to set stimulation parameters, and initiate stimulation on-demand via a wireless link to the external device which then sent data and power to the stimulator. Based on results from a prior study on electrical stimulation of the pelvic nerve^21^, we decided to employ a multi-train stimulation approach to void the bladder. At each bihourly stimulation timepoint (i.e. 2, 4, 6, or 8 hours post-surgery (hps)), ten trains of stimulation were delivered with 20s to 30s inter-train gap. At 2, 4, and 6 hps timepoints, stimulation of the pelvic nerve was carried out while taking care to minimize stress to the rat where one of the hind limbs of the subject was gently lifted by the experimenter to aid visual observation of urine output from the urethral meatus. While visual inspection is not a reliable indicator of volume, we chose this method to confirm voiding as it minimized discomfort and did not require the animal to be removed from their home cage. At the 8 hps timepoint, stimulation was performed while gently restraining the rat in order to collect urine output via an Eppendorf tube placed just beneath the urethral meatus for quantification of urine volume using a precision weighing scale.

### 2.4 Unsuccessful pelvic nerve stimulation

For 2 subjects, the delivery of electrical stimulation did not produce any voiding effects, and the post-mortem analysis revealed that the electrodes had migrated away from the pelvic nerve. One subject was anesthetized at 8 hps, and had its bladder analysed for size, spontaneous activity, and capacity. The other subject was kept as for manual expression at 8-12 hours interval (i.e. two or three times daily), which was a typical bladder management protocol for SCI rats, and subsequently anaesthetized at 28 hps for bladder analysis.

### 2.5 Assessment of bladder size, spontaneous activity and capacity

During the terminal anesthetized (1.5% isoflurane) experiment, the bladder was exposed via abdominal incisions. Bladder size was quantified in terms of width (horizontal x-axis) and height (vertical y-axis) using a measurement ruler. For the analysis of spontaneous contractions of the bladder without catherization, brief videos of the exposed bladder were taken at 30 Hz, and subsequent analyses were performed using custom MATLAB software. 200 frames of each video (see Supporting Information), converted to grayscale, were extracted for analysis. Drift analysis was performed to check for respiratory movement artefacts and a customized motion correction algorithm (MATLAB) was applied to reduce movement artefacts across the 200 frames. Next, a ROI of the bladder was selected for pixel analysis. The very edges of the bladder and areas on the bladder with bright saturated pixels were also excluded from further analysis to eliminate artefacts. Standard deviation and overall mean values of the pixels within the ROI were calculated to obtain the coefficient of variation (CV) over the 200 frames to detect areas within the ROI showing spontaneous contractions (i.e. pixels with higher CV values). The bladder capacities of Rat 1 and Rat 2 (Figure 3) were calculated by adding the volume of urine voided by the stimulation and the residual urine volumes. Bladder capacities of Rat 3 and Rat 4 (Supplementary Figure 1) were quantified from the urine collected by manually squeezing the bladders after the bladders were examined for size and spontaneous contractions.

### 2.6 Recording of pelvic nerve ENG, EUS EMG and intravesical pressure

In a separate group of subjects (n = 7 spinal cord intact, and n = 2 complete spinal cord transection), each rat was anesthetized with a mixture of ketamine (37.5 mg/ml) and xylazine (5mg/ml) intraperitoneally. For 2 subjects, the animal was first placed in a prone position to perform a complete spinal cord transection at T8, similar to that described in Section 2.1, and then placed in a supine position for further surgery. The surgical procedures to expose the bladder, pelvic nerve, and external urethral sphincter (EUS) were as described previously^9^. Platinum iridium hook electrodes were implanted unilaterally on the pelvic nerve, and secured with Kwik-Sil for extraneural recording of pelvic nerve electroneurogram (ENG). EUS electromyography (EMG) was recorded using a pair of fine stainless steel wire. For recording using commercial benchtop systems, both ENG and EMG signals were amplified by using the Intan preamplifier 2216, and acquired at either 20 or 30 kHz with the Intan RHD2000 (Intan Technologies) with a 50 Hz notch filter. Intravesical bladder pressure was recorded using a saline-filled catheter (C30PU-RCA1302, Instech Laboratories Inc) connected to a pressure sensor and infusion pump as described previously^9^. Pressure data was also acquired via the Intan RH2000 system, and then low-pass filtered at 30 Hz. Changes in intravesical pressure were referenced to the minimum pressure values obtained within each recording session. Recording sessions lasted from a few minutes to more than 10 minutes. For one spinal cord intact subject, ENG and EMG were also recorded at 42.9 kHz using our custom-built implantable neural amplifier^22^. All ENG and EMG data were bandpass filtered between 300 and 3000 Hz, and between 20 and 500 Hz, respectively, and further analysed off-line using custom MATLAB software.

### 2.7 Estimation of intravesical pressure from pelvic nerve ENG

Filtered pelvic nerve ENG was first rectified and then integrated over a moving 2 s window. The integrated ENG signal was then further smoothed using the conv function (mask window = 3) in MATLAB, and divided by the its maximum value within the same recording session to obtained the normalized (value between 0 and 1) processed ENG signal. In order to obtain the estimated intravesical pressure changes in mmHg, the processed ENG signal was then multiplied by the maximum increase in intravesical pressure obtained from the same recording session. Measured changes in intravesical pressure were downsampled prior to the calculation of the pairwise correlation coefficient between the measured and estimated intravesical pressure using Spearman’s rank order correlation in MATLAB.

## 3 Results

### 3.1 On-demand stimulation of pelvic nerve in awake SCI rats

To investigate the effects of pelvic nerve stimulation after a SCI, a total of 6 subjects underwent the SCI surgery and electrode implantation on the pelvic nerve. However, 2 rats were excluded from further experiments due to poor recovery from the surgery. Out of the remaining 4 rats, 2 rats showed successful pelvic nerve stimulation (Figure 1 and 2), while the other 2 rats did not due to electrode migration (verified upon terminal experiments), and were subsequently treated as a separate set of animals that underwent manual expression every 8 – 12 hours. As schematized in Figure 1A, we implanted a flexible neural clip electrode (Figure 1C) onto the left pelvic nerve (Figure 1B), and used a custom-built wireless implantable stimulator (Figure 1D and 1E) to deliver electrical stimulation to the pelvic nerve via the head-plug in the awake SCI rat. We then stimulated the subjects bihourly post-surgery (timeline shown in Figure 2A). Figure 2B shows a picture of a subject (Rat 1) during the pelvic nerve stimulation at 6 hours post-surgery (hps) where voiding can be observed from the urethral meatus (indicated by the black arrow). The 2 subjects that showed successful voiding did not display any behaviour that indicated discomfort, nor any vocalization during the stimulation. Across the 2, 4, and 6 hps timepoints, both rats showed variability in the number of drops of urine evoked by stimulation (Figure 2C). At 8 hps, we performed a more thorough characterization of the voiding efficiencies of the pelvic nerve stimulation by measuring the stimulation-evoked amount of voiding and the residual amount of urine in the bladder of both subjects, and found efficiencies of 62.98% in Rat 1 and 32.97% in Rat 2 (Figure 2D).

**Figure 1.**
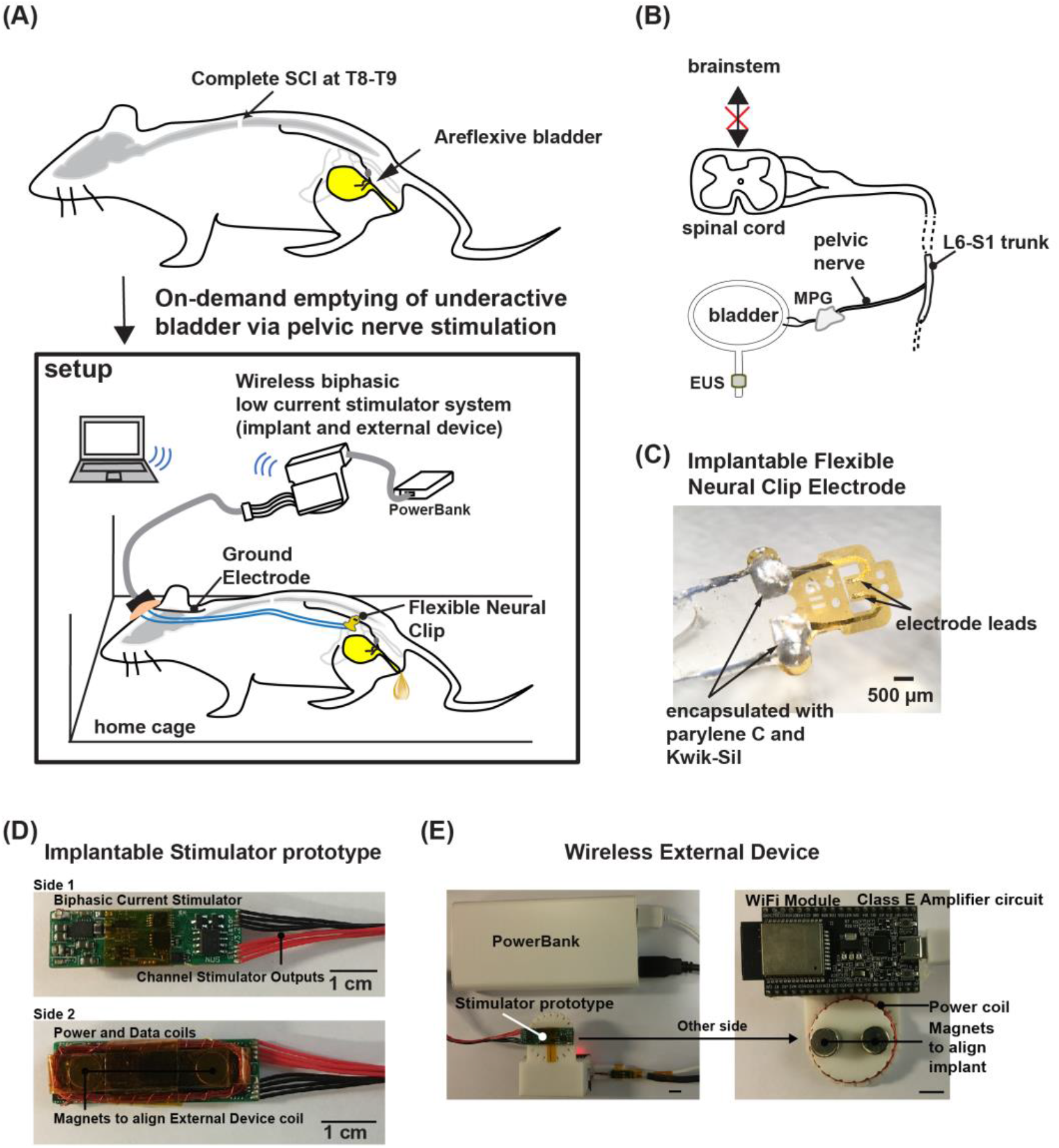
Restoring on-demand voiding of areflexive bladders in SCI rats using pelvic nerve stimulation. (A) Schematic of the setup used to test bi-hourly pelvic nerve stimulation in awake SCI rats in their home cage. (B) The pelvic nerve contains parasympathetic efferent axons that directly drives bladder contraction via postganglionic neurons in the major pelvic ganglion (MPG). The pelvic nerve also contains sensory neurons that convey bladder fullness information back to the brainstem. Spinal cord transection at the T7 level abolishes the micturition reflex, leaving an areflexive bladder. EUS: external urethra sphincter. (C) A flexible neural clip electrode (connected to a headport) was implanted onto pelvic nerves, allowing bipolar stimulation of the pelvic nerves. (D) Photos of the implantable stimulator prototype that was used to deliver biphasic electrical stimulation to the pelvic nerves via the headport attached to the skull. (e) Photos of the wireless external device that provided wireless inductive powering and data to the stimulator prototype. Stimulation parameters were controlled via a customized graphic user interface on the computer, and data were transmitted wirelessly between the external device and computer.

**Figure 2.**
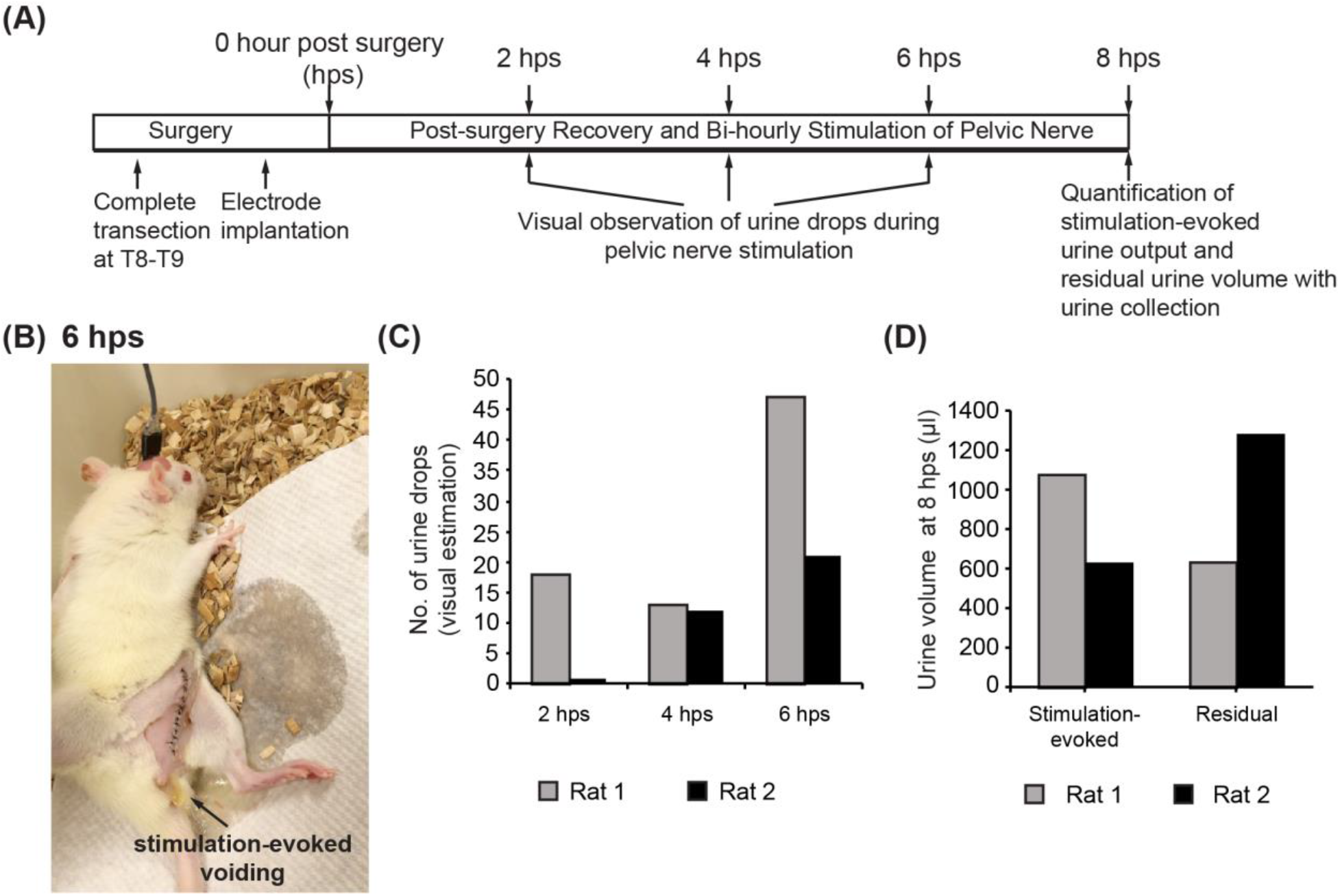
Pelvic nerve stimulation in awake SCI rats evoked bladder voiding. (A) Experimental timeline of the bi-hourly pelvic nerve stimulation for each rat. (B) Photomicrograph of an awake SCI rat in its home cage undergoing the pelvic nerve stimulation 6 hours post-surgery (hps), with urination behaviour observed visually. (C) Number of drops of urine visually observed and estimated during the stimulation at 2 hps, 4 hps, and 6 hps (n = 2). (D) Urine volume collected during the awake pelvic nerve stimulation at 8 hps, and the residual urine volume left in the bladder after stimulation. Residual urine volumes were only measured when the animal was anesthetized, and after the bladder was first examined for size and spontaneous contractions (n = 2).

**Figure 3.**
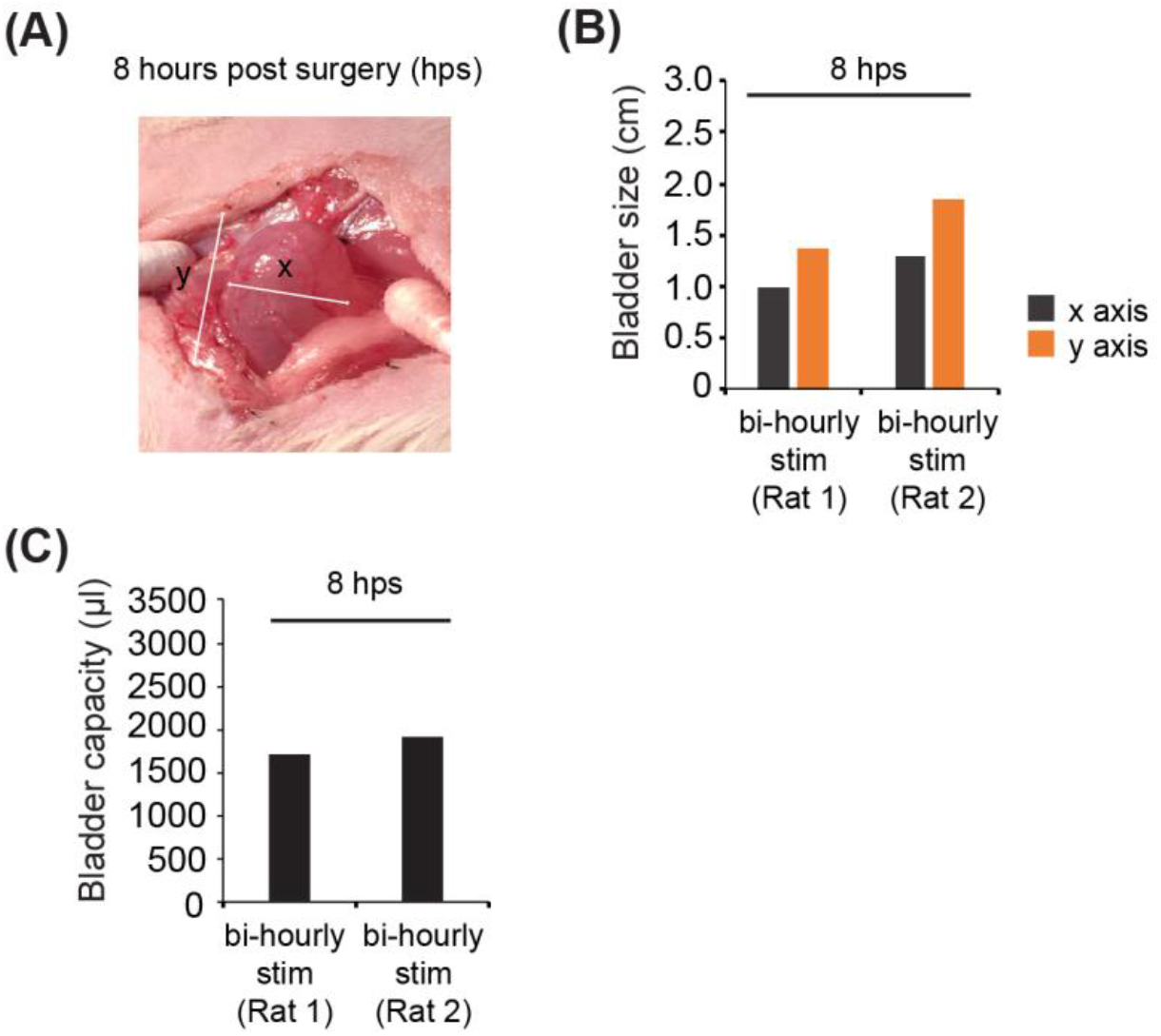
On-demand pelvic nerve stimulation reduced over-distension of bladder. (A) Photomicrograph of a bladder of a SCI rat that underwent successful bi-hourly pelvic nerve stimulation and was examined at 8 hours post surgery (hps). (B) Bladder sizes of the different rats as indicated by the horizontal (x-axis) and vertical (y-axis) lengths of the bladders in (A). (C) Bladder capacities of the pelvic-nerve stimulated rats examined at 8 hps.

### 3.2 Bladder size and capacity of pelvic nerve-stimulated subjects

In order to observe the effect of bi-hourly pelvic nerve stimulation evoked voiding on the bladder, 2 rats were anesthetized for terminal experiments at 8 hps to examine bladder size and capacity (Figure 3, bihourly stimulation *n* = 2). One rat that received no pelvic nerve stimulation within the first 8 hours post SCI surgery was also anesthetized to examine bladder size and capacity (Supplementary Figure 1, Rat 1). One additional non-nerve stimulated subject, which was similarly alert and active after overnight recovery, was subjected to manual expression of the bladder at 8 −12 hours intervals, and anesthetized for a terminal experiment at 28 hps to examine the bladder (Supplementary Figure 1, Rat 2). These 4 rats were the same subjects as described in Section 3.1. Bladder sizes of the subjects that underwent pelvic nerve stimulation had capacities between 1.5 – 2 ml. Subjects that did not receive pelvic nerve stimulation had larger bladder capacities of 2.3 – 3 ml (Supplementary Figure 1) This result was expected as pelvic nerve stimulation every 2 hours produced successful voiding and enabled more frequent voiding, whereas bladders that were only emptied every 8 – 12 hours retained much more urine.

### 3.3 Spontaneous bladder contractions in acute SCI

We also observed the presence of spontaneous bladder contractions (or micromotions) typically observed in the smaller bladders of the subjects that underwent pelvic nerve stimulation. Image analyses of the exposed ventral surface of the bladders (Figure 4A, see Supporting Videos) revealed that bladders from subjects that underwent pelvic nerve stimulation showed spontaneous contraction-induced changes in pixel values over the ventral surface of the bladder. These spontaneous contractions were observed much less in the non-pelvic nerve stimulated subjects (Supplementary Figure 2). These results suggested that frequent emptying to reduce bladder overdistention can preserve spontaneous contractions of the bladder and over-distended bladder walls were compromised in terms of the intrinsic spontaneous excitability within the first 8 to 28 hours of complete SCI, despite manual emptying of the bladder in 8 to 12 hours intervals.

**Figure 4.**
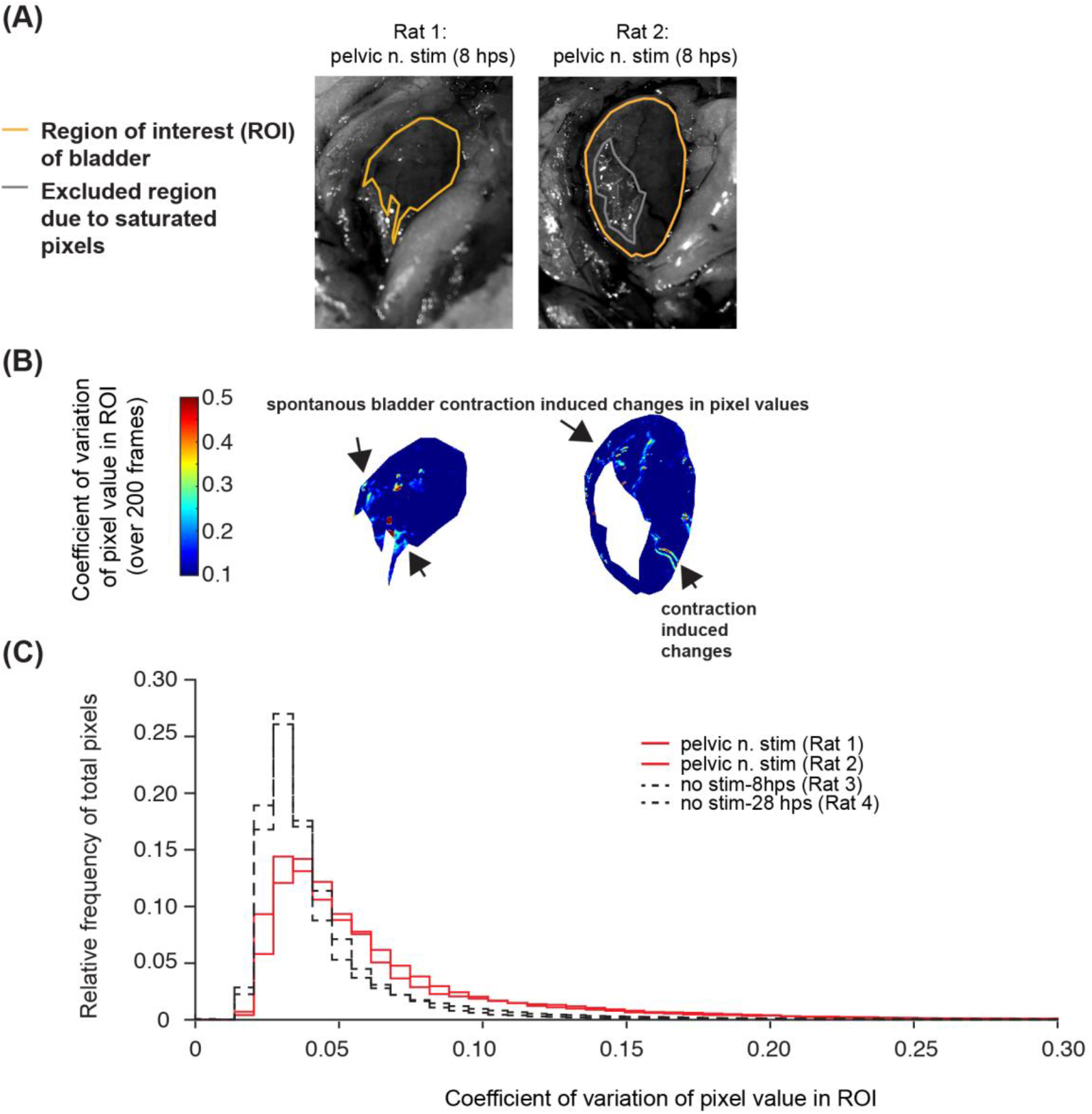
Analysis of bladder spontaneous contractions of SCI rats that underwent pelvic nerve stimulation to evoke voiding. (A) Still gray images of bladders overlayed with regions of interest (ROI) selected for pixel analysis in videos taken from bladders from the 2 rats that underwent pelvic nerve stimulation. Orange lines indicate the boundaries of the ROI region, while the gray lines within the ROI indicate excluded regions that contained oversaturated bright pixels. (B) Colormaps of the various ROIs showing the coefficient of variation (CV) of each pixel value within the ROI over 200 frames in each video (see Supporting Videos). Black arrows indicate pixels with high CV values due to spontaneous contraction movements of the bladder. (C) Relative frequencies of the CV values of pixels in each ROI of the 2 rats. Each colored line indicates a different rat. Data from the 2 non-stimulated rats shown in Supplementary Figure 2 are also plotted here as black dashed lines for visual reference.

### 3.4 Pelvic nerve ENG signals as proxy for intravesical pressure

In order to determine if the pelvic nerve ENG signals obtained at the usual stimulation site could be used to estimate intravesical pressures changes, we performed simultaneous pelvic nerve ENG measurements and intravesical pressure in both spinal intact (n = 7) and spinal transected (n = 2) rats under anaesthesia (Figure 5). Pelvic nerve ENG signals obtained were small in amplitude (typically < 40 μV peak-to-peak) and processed off-line in a series of steps (see Section 2.7) for estimation of intravesical pressures, and compared to the measured pressure changes (Figure 5A). Estimated pressures from ENG were significantly correlated with measured pressures in both spinal intact and spinal transected subjects (an example of each is shown in Figure 5B, R = 0.939 for spinal intact, R = 0.956 for spinal transect, both p < 0.001, Spearman’s pairwise correlation). Although estimated pressures and measured pressures values were not identical, significant (and high) correlation coefficients between estimated and measured values were obtained across all subjects (Figure 5C, all R between 0.827 - 0.956, all p < 0.001, Spearman’s pairwise correlation). EUS EMG signals were also recorded in a subset of animals (4/7 spinal intact animals, an example is shown in Figure 5D), and generally, clear increases in EMG amplitudes were observed in response to larger bladder pressure changes. As a preliminary investigation to see whether an implantable neural recording device could be used to obtain pelvic nerve ENG signals successfully, in one spinal intact subject, we recorded ENG and EMG signals using both the commercial benchtop system (Figure 5D), and a custom-built implantable neural amplifier^22^(Figure 5E) during different recording sessions within the same day. Patterns in ENG and EMG signals obtained from the implantable neural amplifier were comparable to those obtained from the commercial system (Figure 5D and 5E). During the recording session with the implantable system, the actual intravesical pressures were not recorded. Using the same processing steps for the pelvic nerve ENG recorded using the commercial system, we estimated changes in intravesical pressure for that recording session based on the ENG obtained from the implantable system (Figure 5E), and found them to be qualitatively similar.

**Figure 5.**
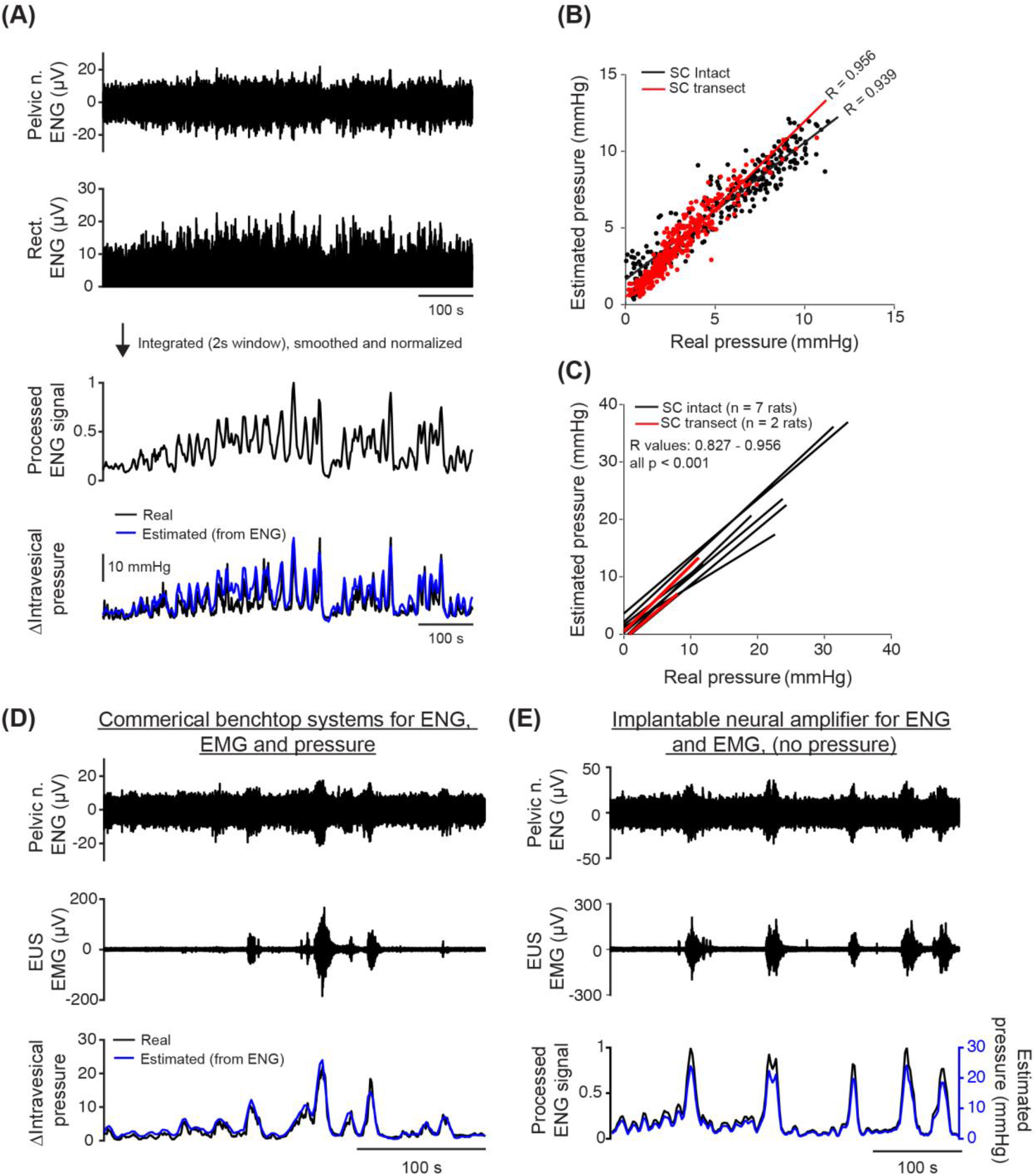
Pelvic nerve signals serve as a proxy for bladder pressure. (A) Pelvic nerve electroneurogram (ENG) recording with simultaneous intravesical pressure recordings in a spinal cord intact rat using commercial benchtop systems. ENG signals were processed in a sequence of steps (rectification (shown in second plot from the top), integration, smoothing and, normalization (shown in third plot from the top)) for estimation of intravesical pressure. Estimated changes in intravesical pressure (blue line, bottom panel) was obtained by multiplying the normalized processed ENG signal with the maximum value of the measured pressure (black line, bottom panel). (B) Examples of correlation between estimated pressure versus measured pressure plots for a spinal cord intact rat (SC intact, black dots and line), and a spinal cord transected rat (SC transect, red dots and line). R values indicate Spearman’s rho. Measured pressure was downsampled to the same sampling rate as the processed ENG signals. Lines indicate linear fits. (C) Summary of estimated pressure versus measured pressure across 9 subjects (n = 7 for SC intact; n = 2 for SC transect). All R values were between 0.827 to 0.956, and p < 0.001 (Spearman’s correlation). (D) Example of simultaneous pelvic nerve ENG recording, external urethral sphincter (EUS) electromyography (EMG) recording, and intravesical pressure recordings using a commercial benchtop system in a SC intact subject. (E) The same subject in (D) was also subsequently recorded for pelvic nerve ENG and EUS EMG using an in-house neural amplifier with no pressure recording capability. Bottom panel shows the processed ENG signal obtained from the neural amplifier (plotted in black), and the estimated pressure (plotted in blue) based on the maximum measured pressure known in (D).

## 4 Discussion

In this preliminary study, we showed that on-demand bi-hourly pelvic nerve stimulation in awake SCI rats using our parameters can evoked successful voiding, and potentially provide an alternative to bladder emptying via manual expression of the bladder (see Figure 1 and 2). Pelvic nerve stimulation reduced bladder overdistension (see Figure 3), and preserved spontaneous bladder contractions (see Figure 4). Based on our previous work^9^, unilateral pelvic nerve stimulation generated changes in intravesical pressures that saturated around 20 mmHg in SD rats, which was similar or even smaller than intravesical pressure changes reported in spontaneous voiding evoked by bladder filling in other studies^23,24^ (peak change of 28 mmHg or more). The typical 8 – 12 hours interval manual expression protocol, which required more physical restraint and handling of the rat, with no potential for remote control, resulted in a more distended bladder. Although the bladder capacity of the all SCI rats examined in this study exceeded the typical rat bladder capacity of about 1 ml^24^, the bladder capacity of the subjects that had its bladder emptied via manual expression were much higher, with a maximum of 3 ml at 28 hps, indicating severe distension. Increasing the frequency of manual expression to bi-hourly may also improve the bladder outcomes, however this hypothesis remains untested in our preliminary study. Thus, an advantage of the pelvic nerve stimulation using implantable devices was that it required less restraint of the subjects than manual expression, thus allowing more frequent emptying.

The variability in the voiding output evoked by nerve stimulation between the subjects (see Figure 3) may be due to differences in the electrode-tissue interface during implantation, or individual variability in the innervation of the bladder by the pelvic nerve. As the complete SCI was performed at the thoracic level above L6-S1, 20 Hz pelvic nerve stimulation alone, in the absence of dorsal root rhizotomy or 20 kHz frequency block, could still evoke reflexive EUS EMG activity, causing detrusor-sphincter dyssynergia^9^, and thus resulting in suboptimal voiding efficiency (32.97% and 62.98%) in our experiments. In order to reduce stimulation-evoked detrusor-sphincter dyssynergia, a combination of low frequency stimulation (10 - 20 Hz) and 20 kHz high frequency nerve block at the pelvic nerve would be required^9^, but these will require further optimization of our implantable electrode and nerve stimulator prototype. Alternatively, a previously published study suggested that voiding efficiency using low frequency stimulation alone could be improved by using an adaptive closed-loop stimulation paradigm instead of a fixed stimulation paradigm^21^. However, this will also require the additional implantation of a bladder pressure sensor.

The physiological role of spontaneous contractions of the bladder is not well-understood, but it has been proposed to be important for maintaining the bladder tone and the dynamic elasticity for efficient voiding^18^. Non-voiding spontaneous bladder contractions are present *in vivo* in bladders across many species (mice, rats, pigs, rabbits, humans, etc.) during the storage phase^25,26^. Our results in this study indicate that bladder emptying using pelvic nerve stimulation in SCI rats, compared to the manual expression technique, resulted in more spontaneous bladder contractions (Figure 4). The spontaneous bladder contractions observed in the pelvic-nerve stimulated subjects were unlikely due to the development of overactivity in the bladder, where bladder capacity is typically decreased^27^. Whether overdistention of the bladder in the early phase of SCI contributes to causing longer-term spontaneous bladder contractions to become greater in amplitude but with reduced frequency^28,29^ remains to be tested. Bladder wall stress has been identified as the primary cause for growth toward hypertrophy^30^, and thus should be avoided as much as possible during bladder management. Gross overdistention of the bladder wall can lead to ischaemia and disruption of the mucosal layer^3^, which may alter the physical and chemical interactions between the detrusor layer and the mucosal layer where suburothelial interstitial cells reside. It has been reported that the removal of the mucosal layer reduced spontaneous contractions of detrusor cells^31^. Overall, these findings indicate the importance for shorter bladder emptying intervals in rats with SCI, and a possible advantage of neural-driven contractions over mechanical or passive emptying methods to better preserve bladder function.

While on-demand pelvic nerve stimulation at prestipulated timepoints can help to empty the bladder, available on-line information about the fullness of the bladder could enable more flexible on-demand scheduling, and more effective delivery of stimulation under a closed-loop neuromodulation scheme^21^. Other than implanting devices that detect bladder size or pressure changes directly^32^, previous studies have shown that it was possible to correlate intravesical pressures with *in vivo* neural recordings obtained from the pelvic nerve^33–35^, pudendal nerve^36^, dorsal root ganglion^37^, and lumbar/sacral roots^38–41^ in various species (e.g. mice, rats, cats, and pigs). In our study, we observed changes in the pelvic nerve ENG that were highly correlated with changes in intravesical pressure in both spinal intact and acutely spinalized subjects (Figure 5). Although extraneural recordings tend to suffer from low signal-to-noise level, and are confounded by the lack of selectivity due to multiunit activity derived from different nerve fibers, our results suggest that extraneural pelvic nerve ENG could be used as a proxy for bladder pressure in spinal cord injury, at least in the early phase. Single unit recordings in L6 dorsal roots in a previous study had revealed that both mechanosensitive Aδ fibers and nociceptive C fibers, which are also present in pelvic nerve can encode information related to bladder-fullness in their firing rate^42^. Despite the potential existence of confounding parasympathetic efferent signals, and cutaneous-related sensory signals^43^ in the pelvic nerve, the significant high correlation between extraneural ENG signals with bladder pressures is likely due to the abundance of afferent fibers in the pelvic nerve, and the inherent causal link between parasympathetic efferent signals and micturition contractions that increase intravesical pressure. Nonetheless, the reconstruction of intravesical pressure traces based on extraneural pelvic nerve ENG still warrants further testing in the awake conscious state in chronic SCI rats to verify its usefulness as part of a closed-loop pelvic nerve-focused neuromodulation strategy for bladder management.

As a preliminary step toward chronic nerve recording, we also performed pelvic nerve ENG and EUS EMG recordings using an implantable neural amplifier^22^ (Figure 5E). The nerve and muscle signals obtained from the implantable neural amplifier were qualitatively similar to those recorded using commercial benchtop system, which allowed us to use the same signal processing steps to estimate intravesical pressure changes. Future work will include more validation experiments, the development of a better way to calibrate the ENG signals to the bladder pressure in awake SCI rats using implantable devices, as well as the longer term stability of the electrode-nerve interface. We acknowledge that the success rate of our experiments can be improved with further optimization of the attachment of the electrode to the nerve. Although there is published evidence indicating that chronic cuff electrodes on pelvic nerve were functionally stable for electrical stimulation for several days post-surgery^44^, it remains unclear if extraneural interfaces can maintain the low impedances required for chronic neural recording. Notably, peripheral nerve recording for predicting bladder pressure remains far from clinical application due to technical and surgical challenges, especially with regards to the electrode-nerve interface as well as the lack of understanding between peripheral nerve signals and bladder volume in human subjects.

## 5 Conclusion

Bi-hourly pelvic nerve stimulation in awake SCI rats can evoked successful voiding, reduced bladder overdistension, and preserved non-voiding spontaneous bladder activity in the areflexive bladder. Extraneural ENG signals recorded from pelvic nerve were also highly correlated with changes in intravesical pressure in both anesthetized spinal intact and SCI rats, enabling the reconstruction of intravesical pressure changes. Results from our study can help improve future designs for implantable devices that combine chronic neural recording and nerve stimulation of the pelvic nerve for closed-loop control of bladder dysfunction after spinal cord injuries.

## Acknowledgements

W.Y.X. Peh, M. Alam, R.U. Ahmed, R. Mogan, G.G.G Gammad designed and performed experiments. S. Lee, L.J. Ong, S. Lu, K.A. Ng, A. Rusly, S. Nag designed, and prepared the devices. W.Y.X. Peh, M. Alam, R.U. Ahmed analyzed the results and wrote the manuscript. Y.P. Zheng and S.C. Yen provided intellectual input, guidance and edited the manuscript. All authors reviewed the manuscript. This work was supported by the GSK Bioelectronic Innovation Challenge (code 100042784), and CRP grant (NRF-CRP10-2012-01) from the National Research Foundation (Singapore), Hong Kong Innovation and Technology Fund (ITS/276/17) and PolyU Research Fund (H-ZG4W).

## Supporting information

1. Supplementary Figure 1

2. Supplementary Figure 2

3. Supplementary video 1: Bi-hourly pelvic nerve stimulation – 8 hps (Rat 1)

4. Supplementary video 2: Bi-hourly pelvic nerve stimulation – 8 hps (Rat 2)

5. Supplementary video 3: No Bi-hourly pelvic nerve stimulation – 8 hps (Rat 3)

6. Supplementary video 4: Manual expression at 8-12 hours interval – 28 hps (Rat 4)

