## Supplementary for "Neuromodulation of pelvic nerve during early phase of spinal cord injury in rats using implantable prototype devices - a preliminary study"

*Corresponding Author

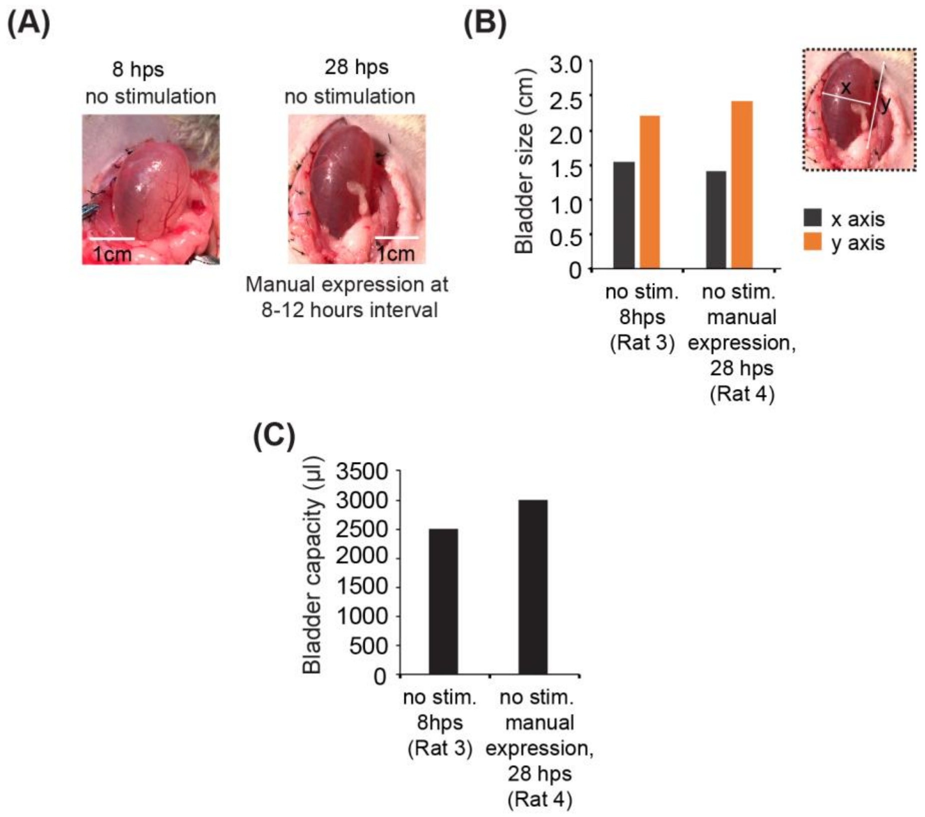


**Supplementary Figure 1. Bladder size and capacity of the non-pelvic nerve stimulated SCI subjects at either 8 or 28 hours post-surgery.** (A) Photomicrographs of the bladders of the non-stimulated rats. One subject was examined at 8 hours post SCI surgery (hps), while another subject was examined at 28 hps with its bladder emptied using manual expression at 8 - 12 hours intervals only. (B) Bladder sizes of the different rats as indicated by the horizontal (x-axis) and vertical (y-axis) lengths of the bladders when examined during terminal anesthetized experiments. (C) Bladder capacities of the rats when examined during terminal anesthetized experiments.

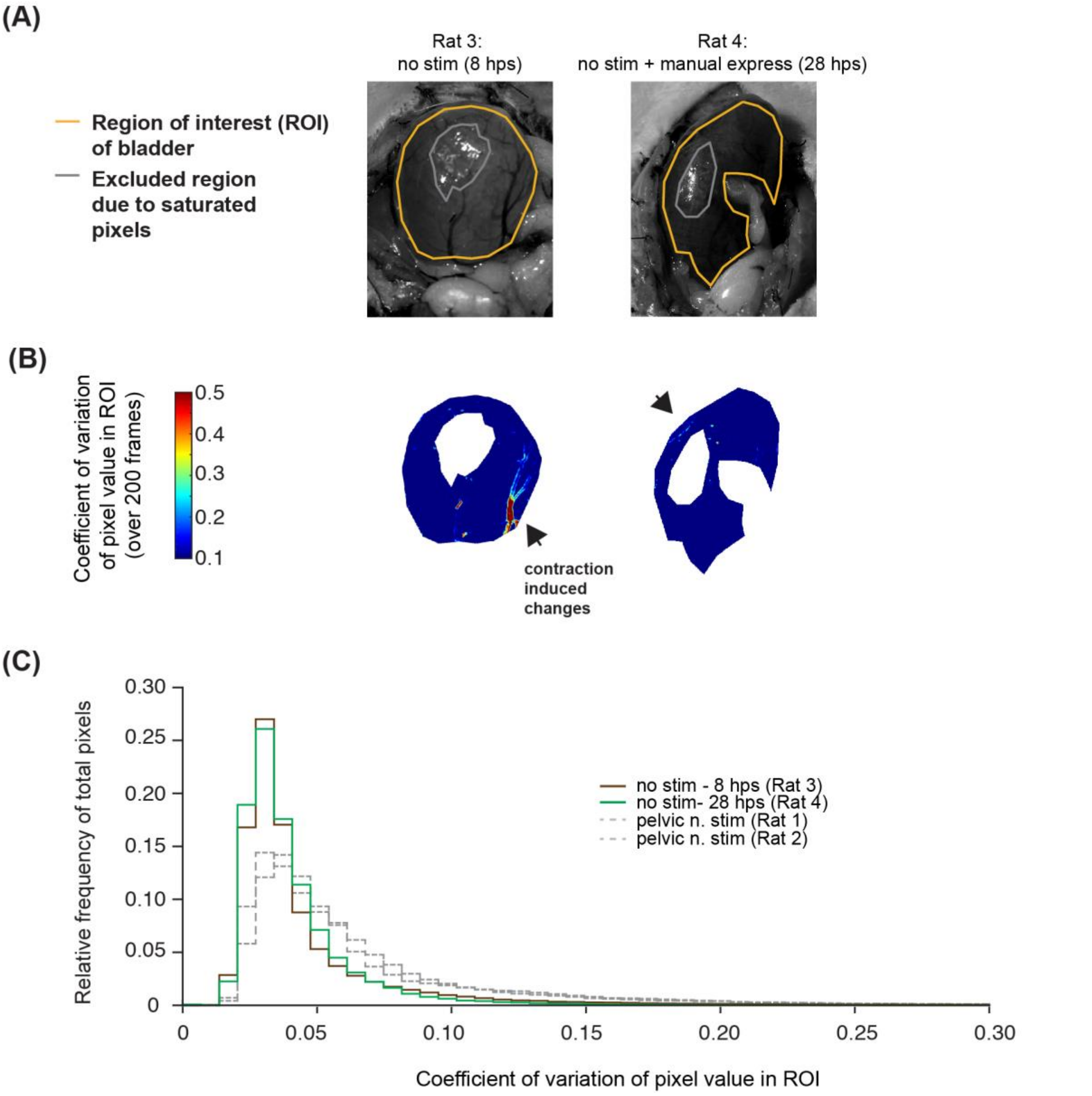


**Supplementary Figure 2. Analysis of bladder spontaneous contractions in non-pelvic nerve stimulated SCI subjects.** (A) Still gray images of bladders overlayed with regions of interest (ROI) selected for pixel analysis in videos taken from bladders. Orange lines indicate the boundaries of the ROI region, while the gray lines within the ROI indicate excluded regions that contained oversaturated bright pixels. (B) Colormaps of the various ROIs showing the coefficient of variation (CV) of each pixel value within the ROI over 200 frames in each video (see Supporting Videos). Black arrows indicate pixels with high CV values due to spontaneous contraction movements of the bladder. (C) Relative frequencies of the CV values of pixels in each ROI of the 2 non-stimulated rats. Each colored line indicates a different rat. Data from the 2 pelvic nerve stimulated rats shown in Figure 4 are also plotted here as gray dashed lines for visual reference.
